## Supplementary Information for "Inference and analysis of cell-cell communication using CellChat"

This file includes the following subsections:

- Supplementary Text
  - Details of method comparisons
  - Evaluation metrics
  - References
- Supplementary Figures 1-8
- Supplementary Table 1

### Supplementary Text

#### Details of method comparisons

**Details of running SingleCellSignalR.** SingleCellSignalR v0.0.1.4[1] was used for inferring intercellular communications of scRNA-seq datasets (https://github.com/SCA-IRCM/SingleCellSignalR_v1). To infer both "autocrine" and "paracrine" signaling, we used “cell_signaling” function with the “int.type” parameter being "autocrine" and the “species” parameter being "mus musculus".

**Details of running iTALK.** iTALK v0.1.0[2] was used for inferring intercellular communications of scRNA-seq datasets (https://github.com/Coolgenome/iTALK). Since iTALK only provides the ligand-receptor database for human, we first map the gene symbols from mouse to human and then run the default iTALK workflow provided in https://github.com/Coolgenome/iTALK/blob/master/example/example_code.r.

**Details of running CellPhoneDB.** CellPhoneDB v2.0.0[3] was used for inferring intercellular communications of scRNA-seq datasets (https://github.com/Teichlab/cellphonedb). Since CellPhoneDB only provides the ligand-receptor database for human, we first map the gene symbols from mouse to human and then run “cellphonedb method statistical_analysis metadata.txt counts.txt --iterations=100 --threads=2”.

#### Evaluation metrics

We evaluated the robustness of inferred interactions from subsampled datasets using three measures, including true positive rate (TPR), false positive rate (FPR) and accuracy (ACC) by comparing the subsampled dataset with the original dataset. They are defined as follows

$$TPR = \frac{TP}{TP + FN}$$

$$FPR = \frac{FP}{TN + FP}$$

$$ACC = \frac{TP + TN}{TP + TN + FP + FN}$$

where TP (true positive) is the number of interactions inferred from the subsampled dataset matched by the interactions inferred from the original dataset, FP (false positive) is the total number of interactions of the subsampled dataset minus TP, TN (true negative) is the number of interactions not in both the subsampled dataset and the original dataset, and FN (false negative) is the number of interaction of the original dataset that are not matched by the subsampled dataset.

### Supplementary Figures

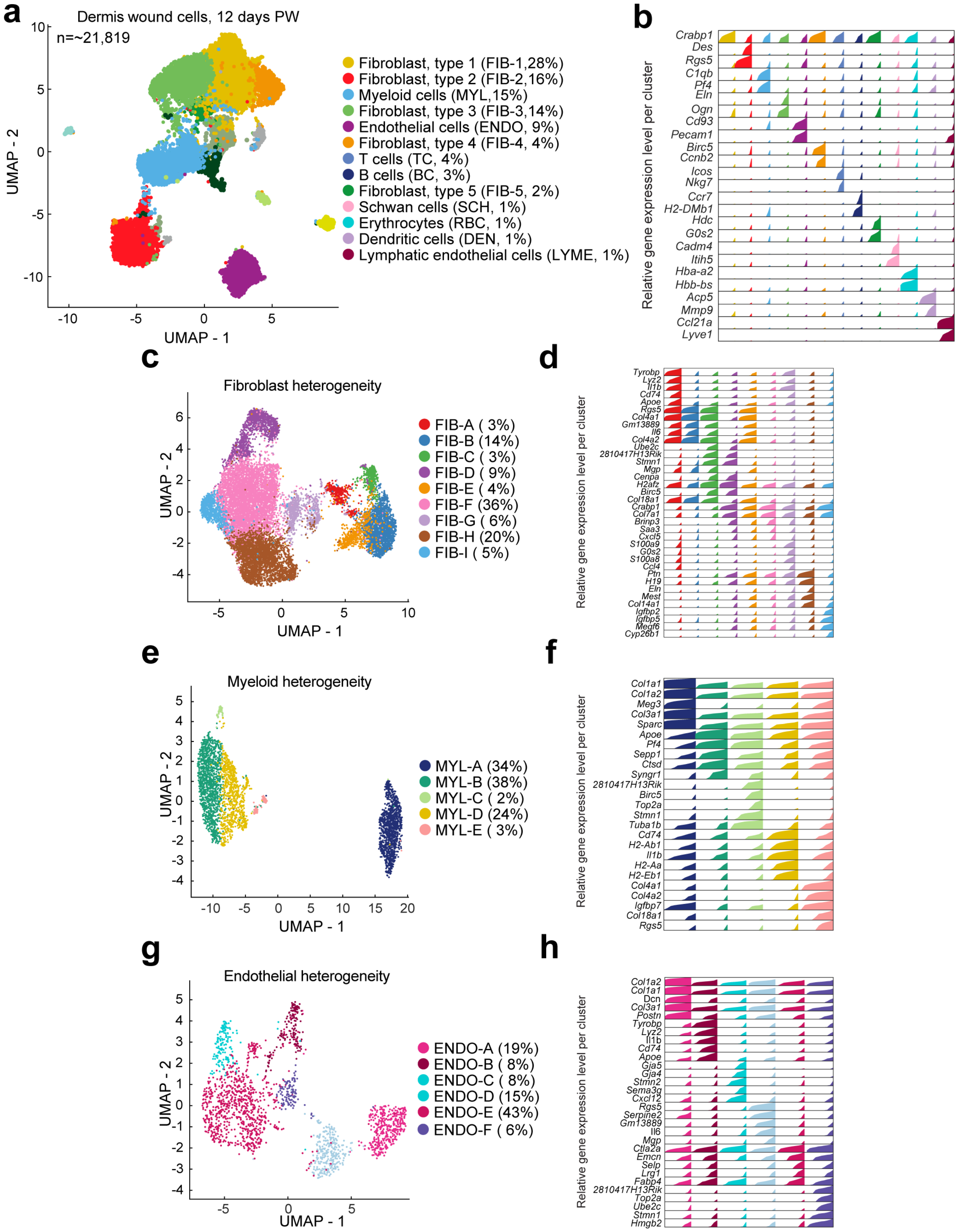

**Supplementary Figure 1: Classification of skin wound cells into groups. (a)** Classification of day 12 skin wound cells into major cell populations. Proportion of cells in each cell group is indicated. **(b)** High-density bar chart showing the distribution of selected marker genes associated with each cell group. Colors correspond to cell populations defined in panel **(a)**. **(c)** Classification of all skin wound fibroblasts into subpopulations. **(d)** Distribution of selected marker genes associated with each fibroblast subpopulation. **(e)** Classification of all skin wound myeloid cells into subpopulations. **(f)** Distribution of selected marker genes associated with each myeloid cell subpopulation. **(g)** Classification of all skin wound endothelial cells into subpopulations. **(h)** The distribution of selected marker genes associated with each endothelial subpopulation.

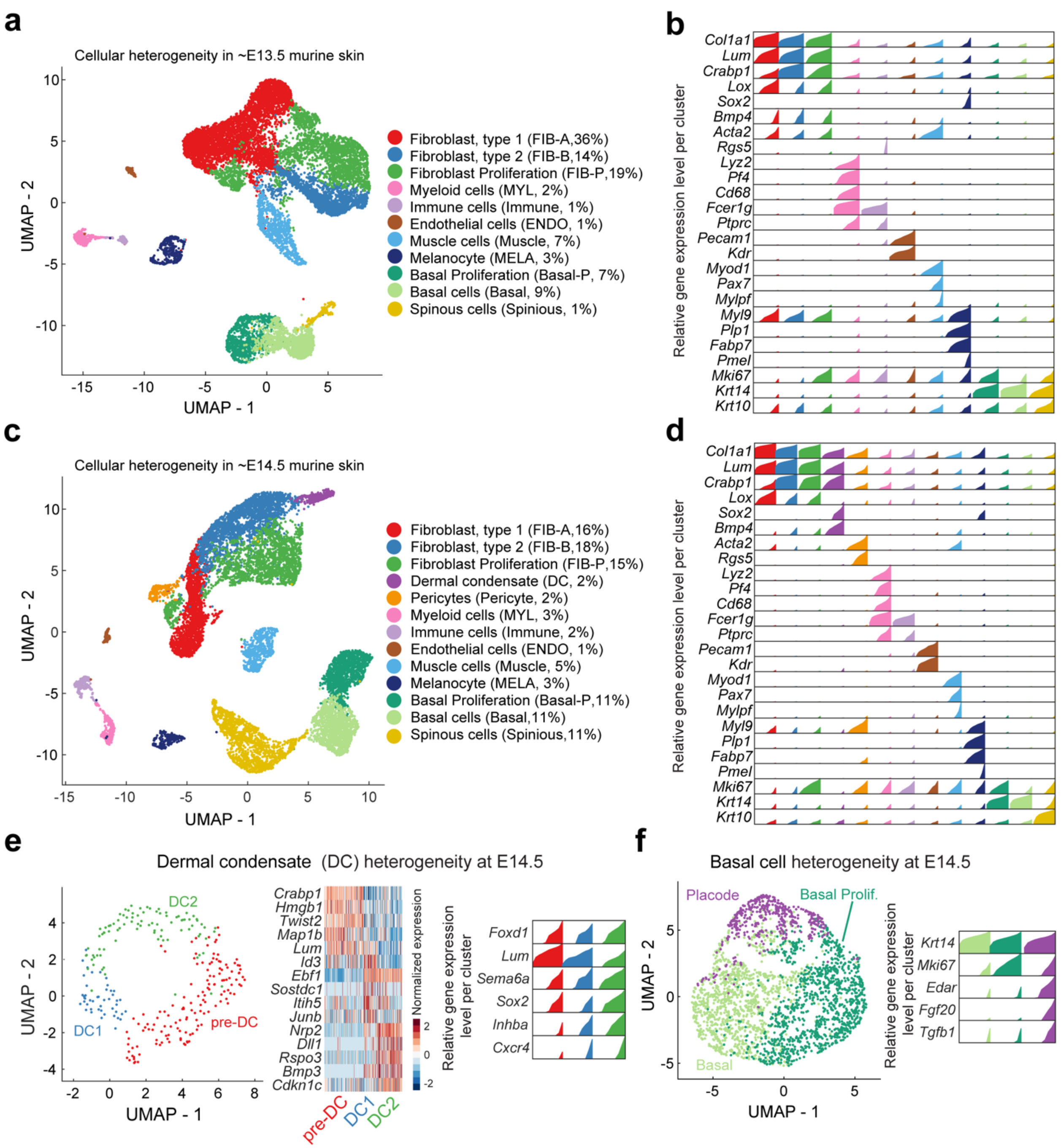


**Supplementary Figure 2: Classification of cells from embryonic skin dataset into groups. (a)** Classification of skin cells from E13.5 into major cell populations. The proportion of cells in each population is indicated. **(b)** High-density bar chart showing the distribution of selected marker genes associated with each cell population from E13.5. **(c)** Classification of skin cells from day E14.5 into major cell populations. Proportion of cells in each population is indicated. **(d)** High-density bar chart showing the distribution of selected marker genes associated with each cell population from E14.5. Colors correspond to the cell populations. **(e)** Classification of dermal condensate cells into subpopulations. The heatmap and the distribution of selected marker genes associated with each subpopulation are shown. **(f)** Left: Classification of basal epidermal cells into subpopulations. Right: The distribution of selected marker genes associated with each subpopulation.

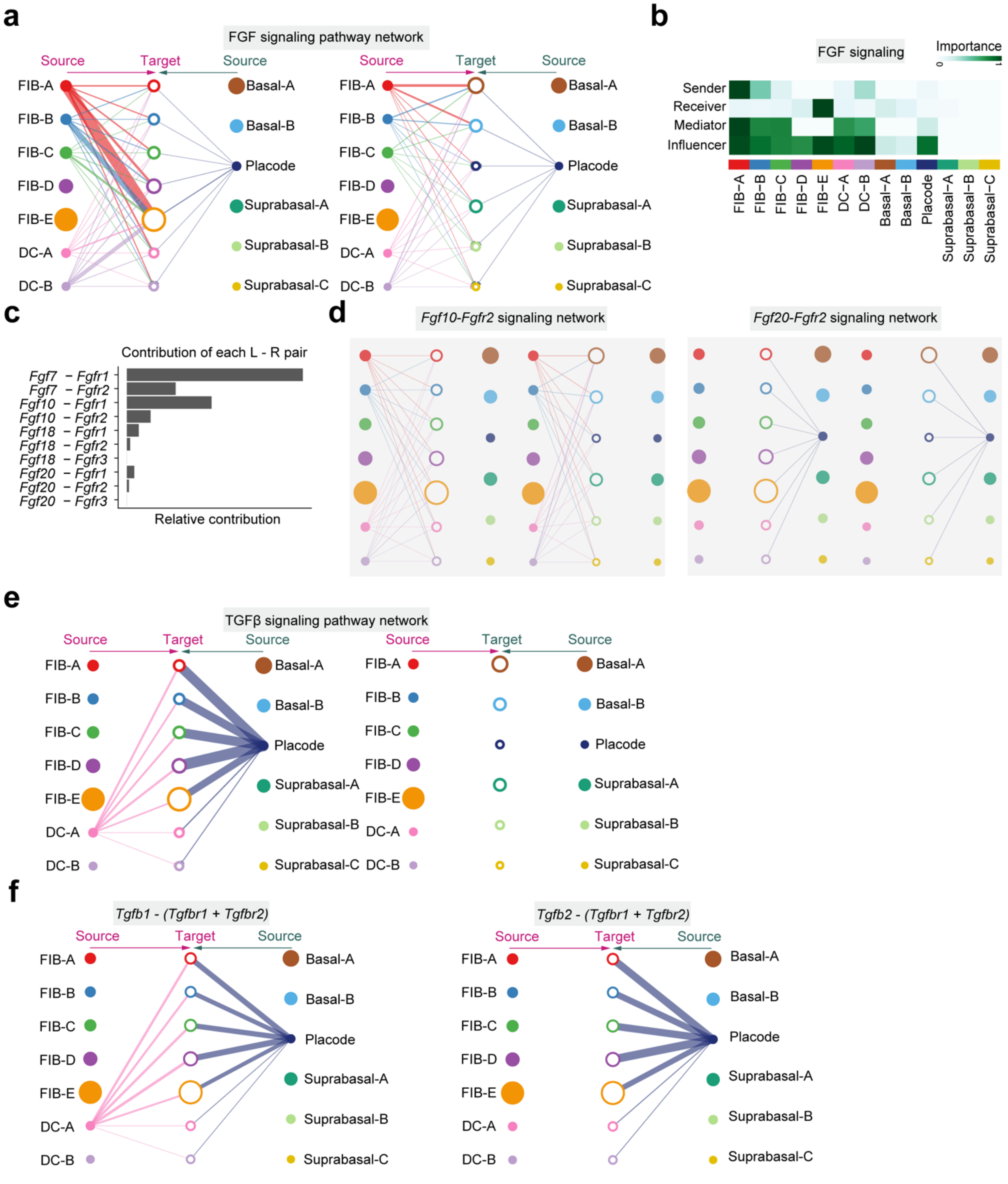

**Supplementary Figure 3: Example intercellular communication networks for continuous cell states along cell pseudotemporal trajectories.** **(a)** Hierarchical plot showing dermal and epidermal communications via FGF signaling. **(b)** Heatmap showing the relative importance of each cell group based on the computed four network centrality measures of FGF signaling. **(c)** Relative contribution of each ligand-receptor pair to the overall communication network of FGF signaling pathway. **(d)** Inferred intercellular communication networks of two ligand-receptor pairs, Fgf10-Fgfr2 and Fgf20-Fgfr2. **(e)** Inferred intercellular communication networks of TGFβ signaling pathway. **(f)** Inferred intercellular communication networks of two ligand-receptor pairs, *Tgfb1 – (Tgfbr1+Tgfbr2)* and *Tgfb2 – (Tgfbr1+Tgfbr2).*

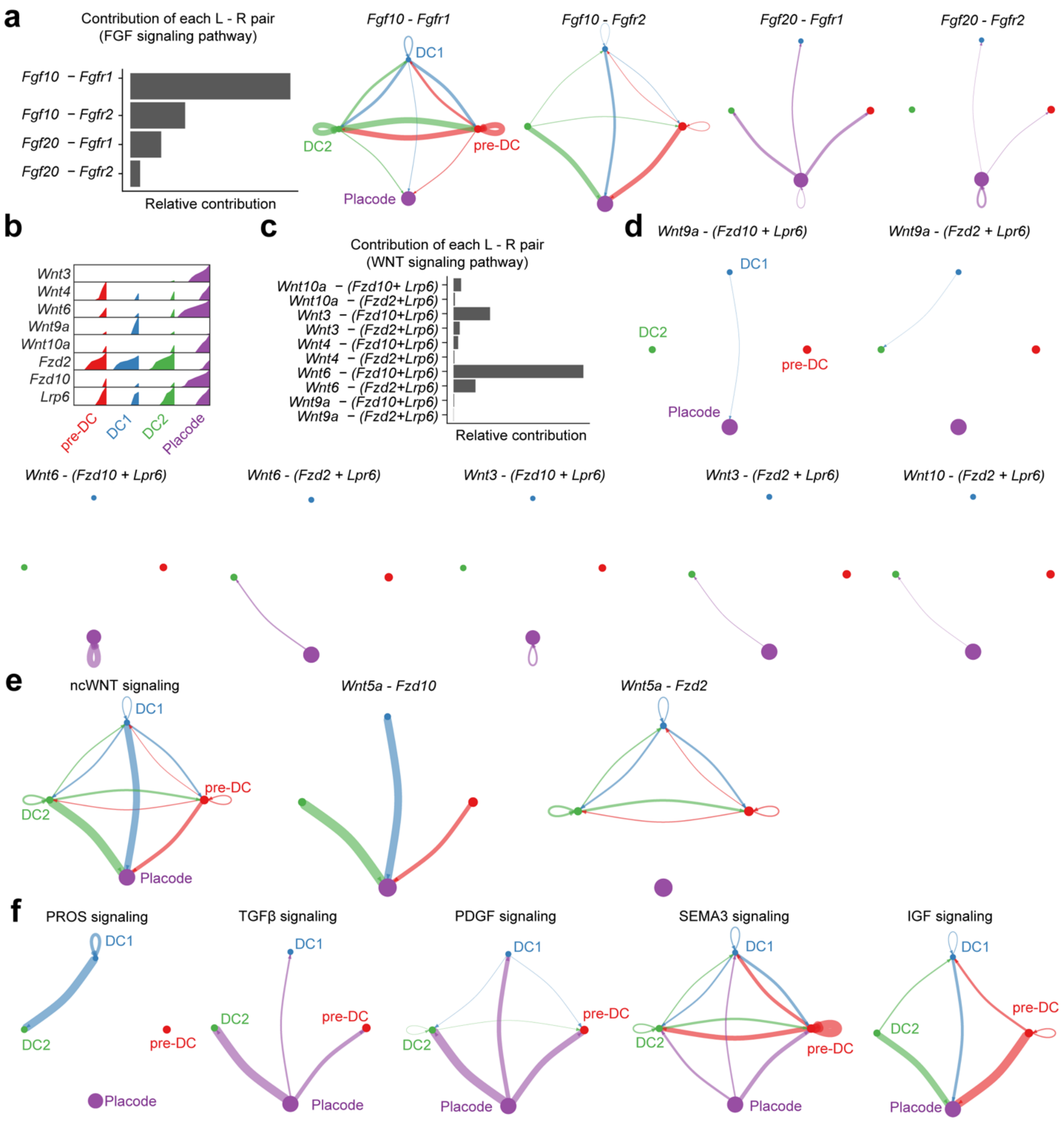


**Supplementary Figure 4: Example intercellular communication networks for spatially colocalized DC and placode cell populations. (a)** Left: Relative contribution of each ligand-receptor pair to the overall communication network of FGF signaling pathway. Right: Inferred intercellular communication network for each ligand-receptor pair associated with FGF signaling pathway. **(b)** The distribution of several WNT pathway signaling genes in each cell group. **(c)** Left: Relative contribution of each ligand-receptor pair to the overall communication network of WNT signaling pathway. Right: Inferred intercellular communication network for some ligand-receptor pairs associated with WNT signaling pathway. **(d)** Inferred intercellular communication network of ncWNT signaling pathway and associated ligand-receptor pairs. **(e)** Inferred intercellular communication networks for PROS, TGFβ, PDGF, SEMA3 and IGF signaling pathways.

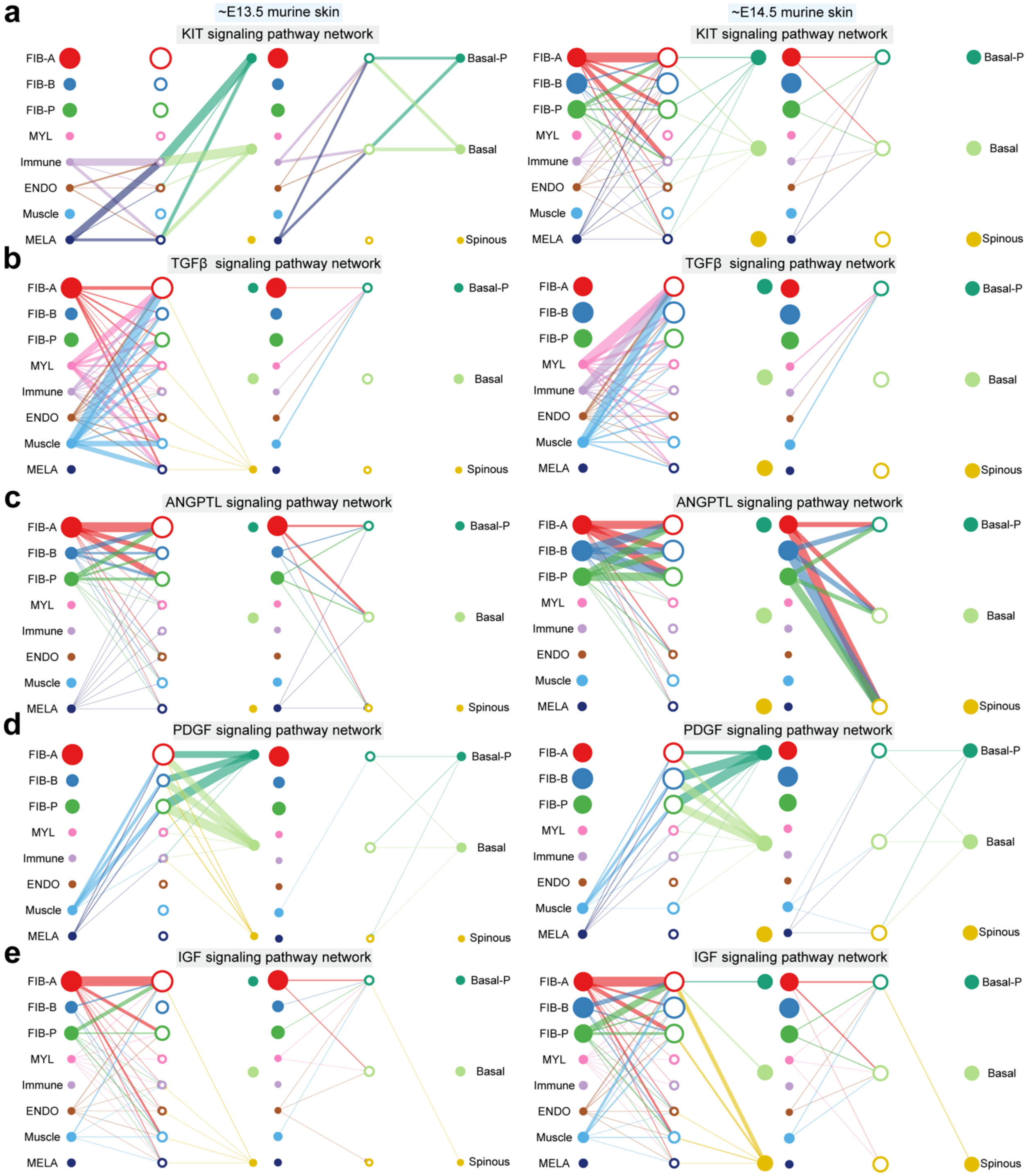


**Supplementary Figure 5: Comparison of example intercellular communication networks between E13.5 and E14.5 embryonic skin. (a-e)** Inferred intercellular communication networks for KIT, TGFβ, ANGPTL, PDGF and IGF signaling pathways at E13.5 and E14.5.

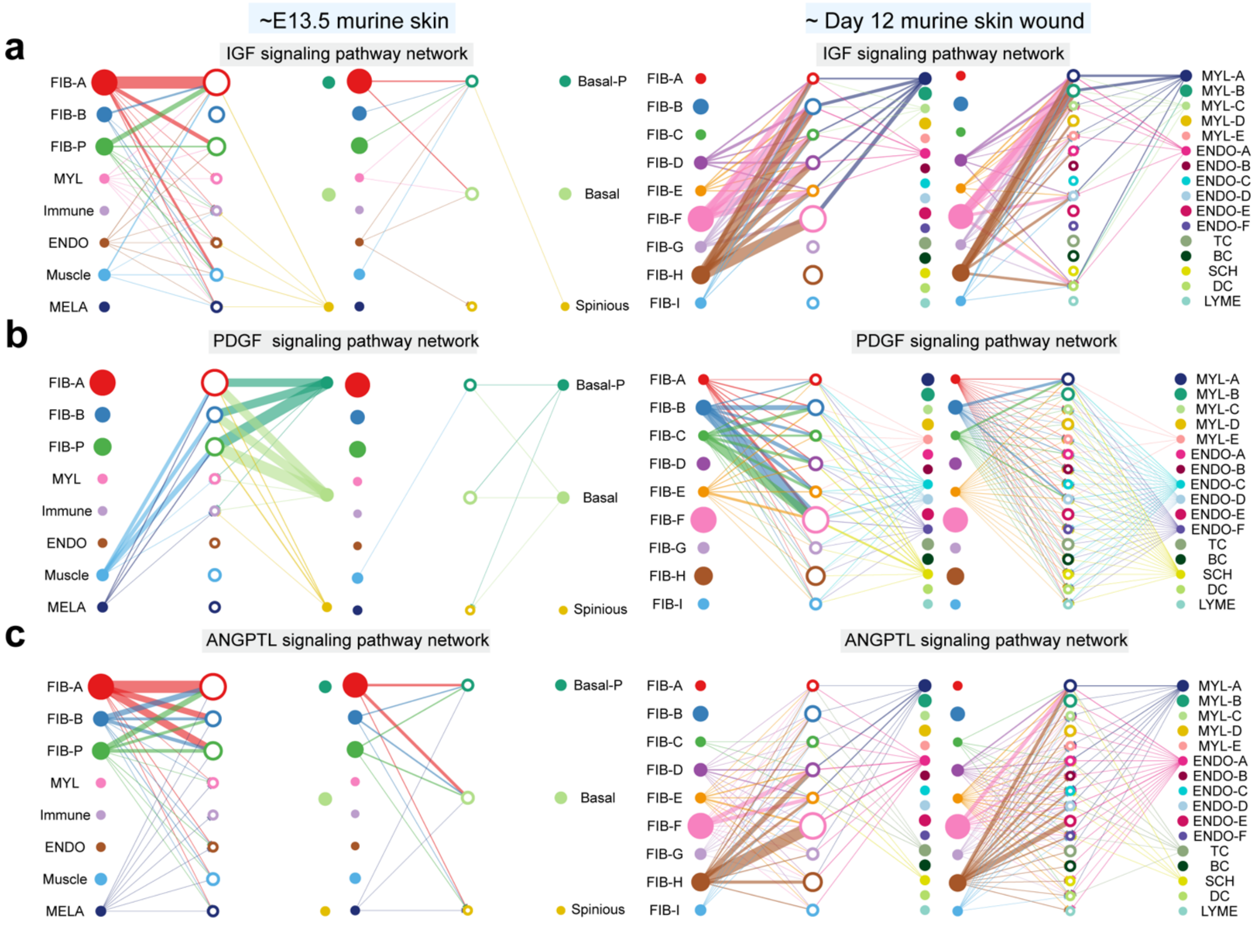


**Supplementary Figure 6: Comparison of example intercellular communication networks between E13.5 embryonic skin and adult skin wound. (a-c)** Inferred intercellular communication networks for IGF, PDGF and ANGPTL signaling pathways in embryonic skin at E13.5 and adult skin wound at day 12.

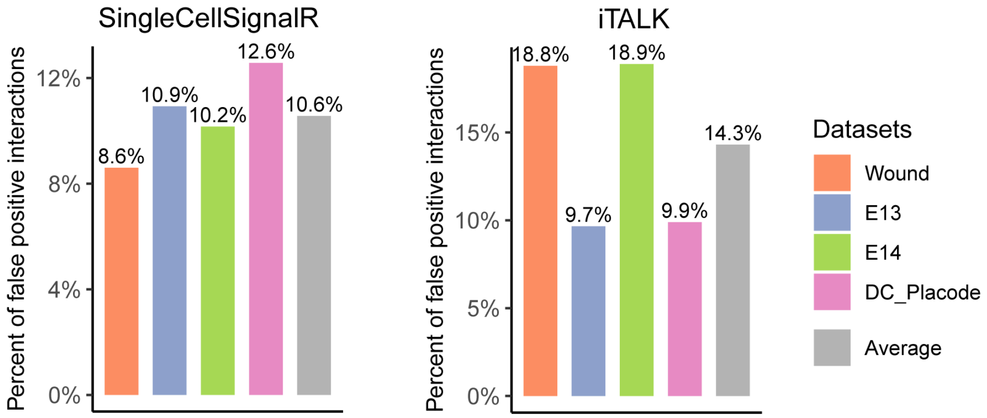


**Supplementary Figure 7: The percentage of false positive interactions caused by incomplete representations of known signaling molecule interactions.** We compute the percentage of false positive interactions inferred by SingleCellSignalR and iTALK. The false positive interactions are defined by the interactions with multi-subunits that are *partially* identified by these tools.

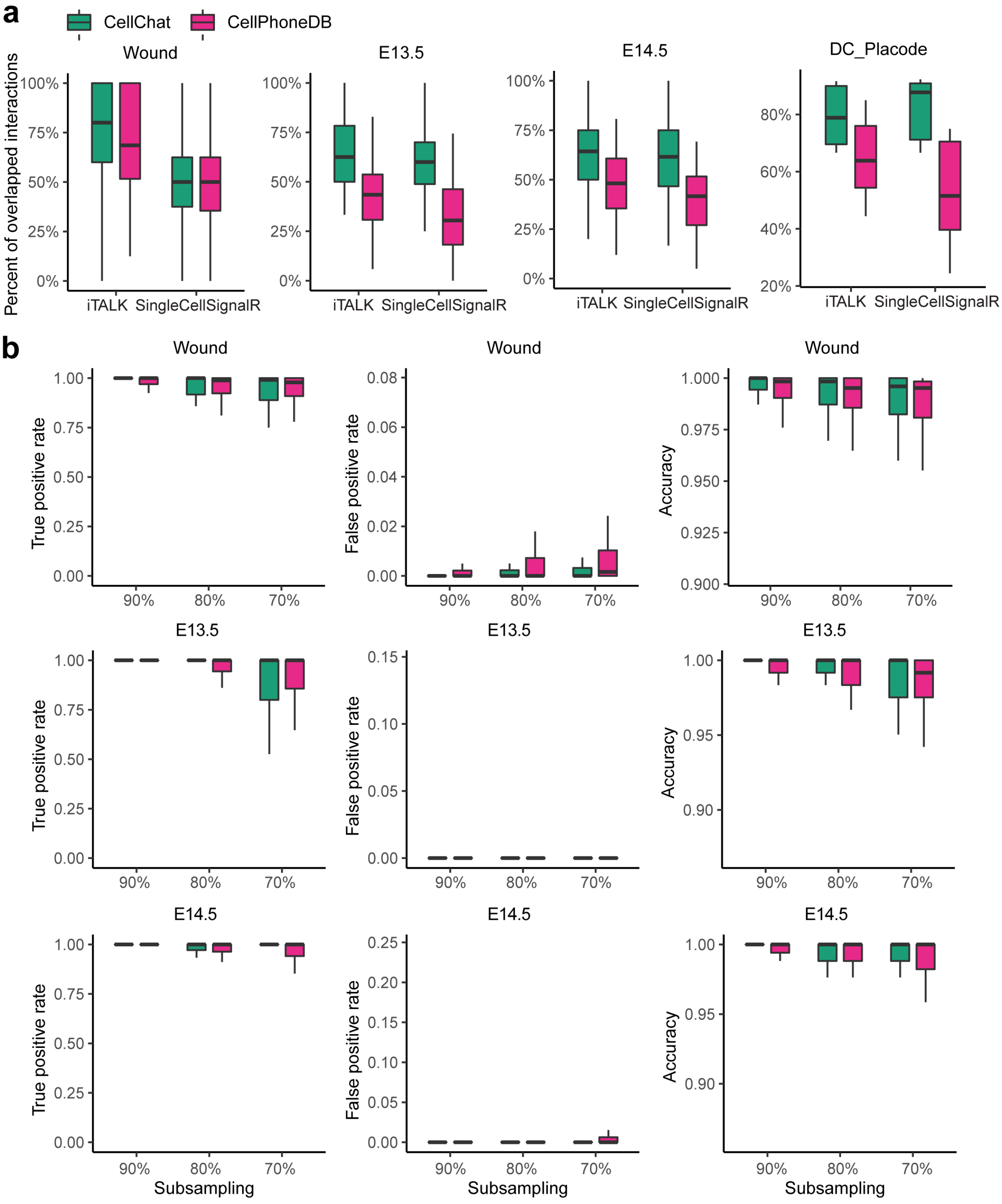


**Supplementary Figure 8: Comparison of ligand-receptor interactions predicted by CellChat and CellPhoneDB. (a)** The percentage of overlapped ligand-receptor interactions between CellChat/CellPhoneDB and other two methods including SingleCellSignalR and iTALK are shown. Each colored bar represents a dataset, and the grey bar indicates the average percentage of overlapped interactions across all datasets. The overlap percentage is defined as the number of overlapped ligand-receptor interactions divided by the minimum number of identified ligand-receptor interactions between two methods. **(b)** Comparison of the robustness of inferred interactions under subsampling (90%, 80%, 70%) of the cells from each dataset. In each case, we compute three measures including true positive rate (TPR), false positive rate (FPR) and accuracy (ACC) by comparing the subsampled dataset with the original dataset.

### Supplementary Table

**Supplementary Table 1. Characterization and comparison of CellChat with other tools for intercellular communication analysis.**

| | | CellChat | SingleCellSignalR | iTALK | CellPhoneDB |
|---|---|---|---|---|---|
| **Database** | Multisubunit structure | Y | N | N | Y |
| | Cofactors | Y | N | N | N |
| | Structured pathways | Y | N | N | N |
| | Number of interactions | 2021 | 3251 | 2648 | 1396 |
| | Species | mouse/human | mouse/human | human | human |
| **Input data** | Preprocessed data matrix | Y | Y | Y | Y |
| | Low-dimensional space | Y | N | N | N |
| | Multiple datasets | Y | N | Y | N |
| **Model** | Methodology | Mass Action Law + Statistical test | Regularized product + Thresholding | DEG analysis | Mean expression + Statistical test |
| | Cell proportion | Y | N | N | N |
| **Systems analysis** | Infer signaling roles of cells | Y | N | N | N |
| | Predict key incoming and outgoing signals | Y | N | N | N |
| | Predict signaling patterns | Y | N | N | N |
| | Classify signaling networks | Y | N | N | N |
| | Identify conserved vs. context-specific signaling | Y | N | Y | N |
| **Visualization** | Hierarchical plot | Y | N | N | N |
| | Circle plot | Y | N | Y | N |
| | Bubble plot | Y | N | N | Y |
| | Alluvial plot | Y | N | N | N |
| **Language** | | R | R | R | Python |
